# Natural killer cell migration in HIV-infected individuals is inhibited by impairment of HIF-1α-mediated glycolysis

**DOI:** 10.1101/2024.05.31.596765

**Authors:** Xiaowen Yu, Jie Zhou, Jie Lei, Hongchi Ge, Zining Zhang, Yajing Fu, Xiaoxu Han, Qinghai Hu, Haibo Ding, Wenqing Geng, Hong Shang, Yongjun Jiang

## Abstract

Natural killer (NK) cells serve as the first line of defense of the immune system and play a crucial role in fighting against HIV infection. The effective function of NK cells is closely related to their migration ability, but the status of NK cell migration in HIV-infected individuals and the regulation mechanism for NK cell migration remains unknown. Here, we found that NK cell migration was significantly impaired in HIV-infected individuals, lower in immune non-responders (INRs) compared with immune responders (IRs), and positively correlated with CD4^+^ T cell counts. Further investigations showed that the decreased NK cell migration in HIV infection was caused by the impairment of glycolysis. Mechanistically, we found that NK cell migration was regulated by HIF-1α pathway, and inhibitory receptor TIGIT restrained HIF-1α expression by inhibiting PI3K/AKT/mTORC1 or ERK signaling pathway, consequently weakening the glycolysis of NK cells in HIV-infected individuals, and ultimately leading to down-regulation of migration. Collectively, we uncovered a mechanism of reduced NK cell migration in HIV infection and provided a new insight for immunotherapy in HIV infection.

**In Brief:** The effective function of NK cells is closely related to its migration ability. The authors show that impaired NK cell migration in HIV-infected individuals is caused by TIGIT inhibiting HIF-1α-mediated glycolysis via PI3K/AKT/mTORC1 or ERK pathway.

## Introduction

Acquired immunodeficiency syndrome (AIDS), an infectious disease caused by human immunodeficiency virus (HIV) infection, can progressively damage the human immune system, induce a series of infections, inflammation, and malignant tumors, and eventually lead to death^1, 2^. Although the mortality rate of HIV patients has been greatly reduced with the widespread use of highly active antiretroviral therapy (HARRT), AIDS is still one of the most serious global public health problems^3^.

Natural killer (NK) cells, an important component of innate immune cells, play a crucial role in antiviral infection and tumors^4, 5^. NK cells are the first line of defense of the immune system, with the functions of cytotoxicity and cytokine secretion^6, 7^, and the effective function of NK cells is related to their migration ability^8^. During HIV infection, it has been reported that chemokines CCL3, CCL4, and CCL5 secreted by NK cells can limit viral replication in vitro^9^. Moreover, NK cells can eliminate HIV-infected cells through antibody-dependent cell-mediated cytotoxicity(ADCC) and the release of perforin, granzyme, and CD107a^10, 11^. Meanwhile, HIV infection resulted in a decrease in the number of NK cells, abnormal distribution of cell subsets, disrupted expression of surface markers, and decreased ability of NK cells to lyse virus-infected cells^11–14^. However, the migration of NK cells in HIV-infected individuals and its relationship with disease progress has yet to be reported.

Chemokines/chemokine receptors axis and cytoskeletal rearrangement exert a pivotal role in the migration of NK cells^15^. The binding of chemokines to their receptors triggers complex signaling cascades that regulate cytoskeletal renewal and induce cell migration^16^. Cytoskeletal rearrangement is conducted by the relative slippage between actin filaments (F-Actin) and myosin^17^. Actin-binding protein, cofilin, modifies actin cytoskeletal architecture by modulating actin polymerization and depolymerization, ultimately promotes cell migration and motility^18–21^. NK cells are known to effectively inhibit tumor cell growth and viral infection, with improved prognosis and increased survival rates in tumor patients related to higher migration ability of NK cells^8, 22^. In addition, it was found that the impaired migration of NK cells to the central nervous system in mice with experimental autoimmune encephalomyelitis (EAE) exacerbated disease severity^23^. Therefore, identifying and understanding the changes and regulation mechanisms of NK cell migration is important in HIV infection.

Accumulating evidences have shown that impaired cell metabolism is the key factor leading to NK cell dysfunction in a series of chronic diseases^24^. It is reported that in mouse NK cells, decreased glycolysis may damage cell proliferation and NK cell cytotoxicity^25, 26^. Additionally, the production of IFN-γ and the expression of granzyme B in NK cells could be inhibited by directly limiting the rate of glycolysis by using 2-deoxy-D-glucose (2-DG)^27, 28^. Cell movement is probably the most energy-consuming cellular activity^29^, but the metabolic requirements of NK cell migration have not been studied.

The present study investigated the migration of NK cells in HIV infection, explored the relationship between NK cell migration and HIV disease progression, and found the possible cause of decreased NK cell migration in HIV-infected individuals. Then the signaling pathway and related mechanisms involved in NK cell migration were demonstrated. Finally, this study identified key molecules that could be potential targets for recovering NK cell migration in HIV infection. Together, our study provided novel insight into the therapeutic strategies for HIV infection.

## Results

### NK cell migration is impaired in HIV-infected individuals and positively correlated with CD4^+^ T cell counts

To study the changes of peripheral blood NK cell migration in HIV infection, we induced NK cells to pass through the microporous membrane by adding chemokine C-X-C motif chemokine ligand-12 (CXCL12) to detect the chemotaxis of NK cells. The results showed that CXCL12-mediated NK cell migration was lower in HIV-infected individuals without ART (HIV ART-), immune responders (IRs) and immune non-responders (INRs) than in HC individuals (Fig 1A). Moreover, CXCL12-mediated migration of NK cells in immune non-responders (INRs) was lower than that in immune responders (IRs), suggesting a potential link between NK cell migration and the immune recovery of HIV. During HIV infection, the absolute count of CD4^+^ T cells is a direct indicator of HIV disease progression and immune recovery with ART treatment, so we further analyzed the correlation between NK cell migration and CD4^+^ T cell counts in HIV-infected individuals. Our analysis showed that NK cell migration in HIV-infected individuals was significantly positively correlated with CD4^+^ T cell counts (Fig 1B).

**Fig 1.**
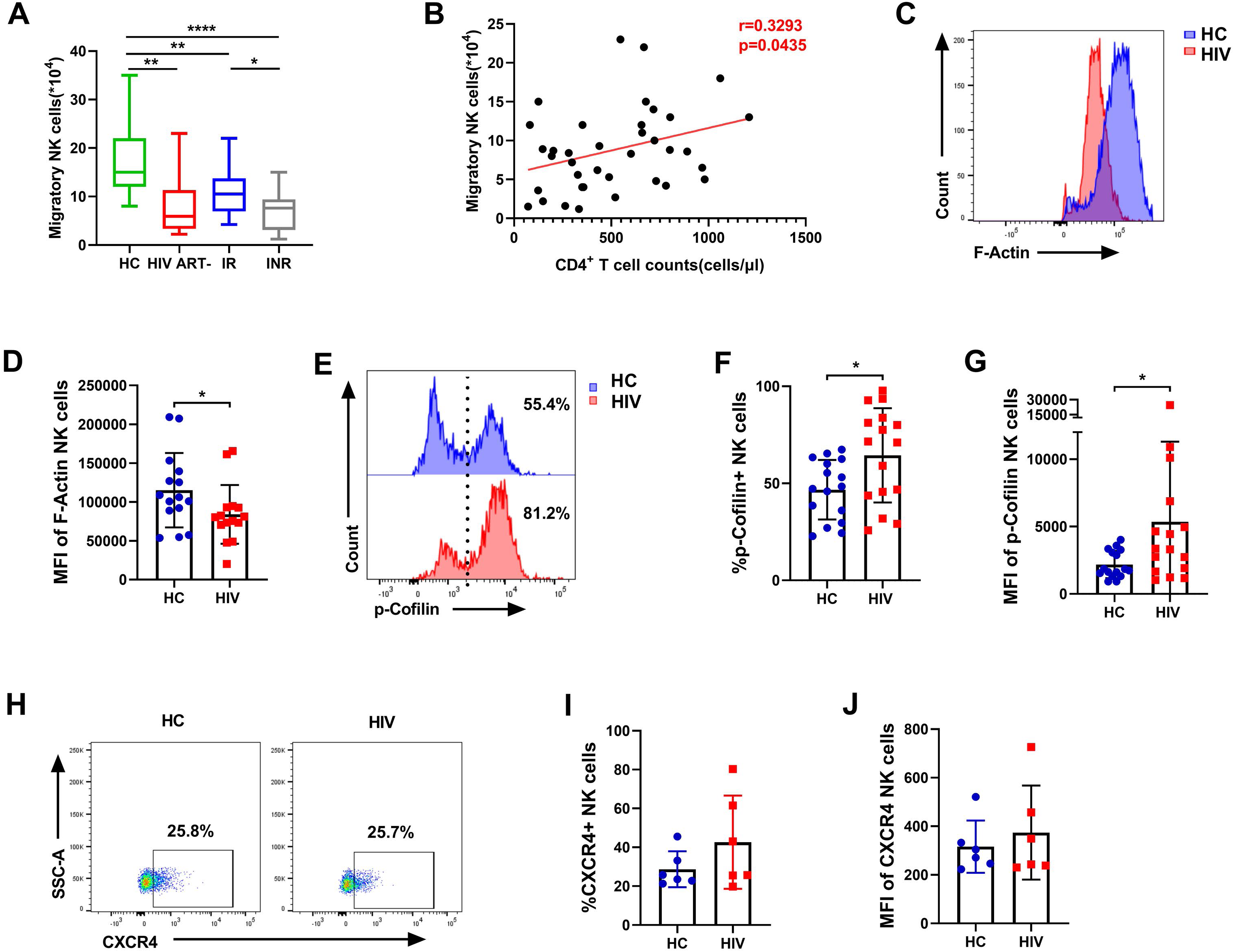
NK cell migration is impaired in HIV infection and positively correlated with CD4^+^ T cell counts. **(A)** CXCL12 (100ng/ml)-triggered migration of NK cells in HC individuals (n = 27), ART-naïve HIV-infected individuals (n = 8), immune responders (IR) (n = 16) and immune non-responders (INR) (n = 14). **(B)** Analysis of the correlation between NK cell migration and absolute CD4^+^ T cell counts (cells/ul) at the same sampling time (n = 38). **(C and D)** The MFI of F-Actin in NK cells from the HC (n = 15) and HIV (n = 15) individuals. **(E - G)** The percentage and MFI of p-cofilin in NK cells from the HC (n = 16) and HIV (n = 16) individuals. **(H - J)** The percentage and MFI of CXCR4 on NK cells from the HC (n = 6) and HIV (n = 6) individuals.

Besides, we detected the levels of F-Actin and cofilin in NK cells, indicators related to cytoskeletal recombination and cell migration. Results showed that the level of F-Actin in HIV-infected individuals was lower than that in HC individuals (Figs 1C and 1D), and the level of Cofilin phosphorylation (inactivate) in HIV-infected individuals was higher than that in HC individuals (Figs 1E-1G).

To explore whether the impaired migration of peripheral blood NK cells in HIV infection is attributed to the alterations in chemokine receptors, we simultaneously detected the expression of CXCR4, CXCL12 chemokine receptor, and the representative flow cytometry plots were shown in Figs 1H. The percentage and mean fluorescence intensity (MFI) of CXCR4 expression on NK cells were comparable between HIV-infected individuals and HC individuals (Fig 1I and 1J), which indicated that the impaired migration of NK cells in HIV infection was not caused by the alterations of chemokine receptors.

### NK cell migration is hindered by down-regulation of glycolysis in HIV-infected individuals

To further explore the factors leading to the impairment of NK cell migration in HIV infection, we analyzed the RNA-seq data of HC individuals and HIV-infected individuals obtained from the data set GSE25669 of GEO database. Gene set-enrichment analysis (GSEA) showed downregulation of the genes related to lymphocyte transendothelial migration and regulation of Actin cytoskeleton in NK cells of HIV-infected individuals compared to HC individuals (Fig 2A). We also found that the mRNA expression of *ACTN1* in HIV-infected individuals decreased significantly, while there was no difference in the mRNA expression of *CXCR4* (Fig 2B), which was consistent with our observations above.

**Fig 2.**
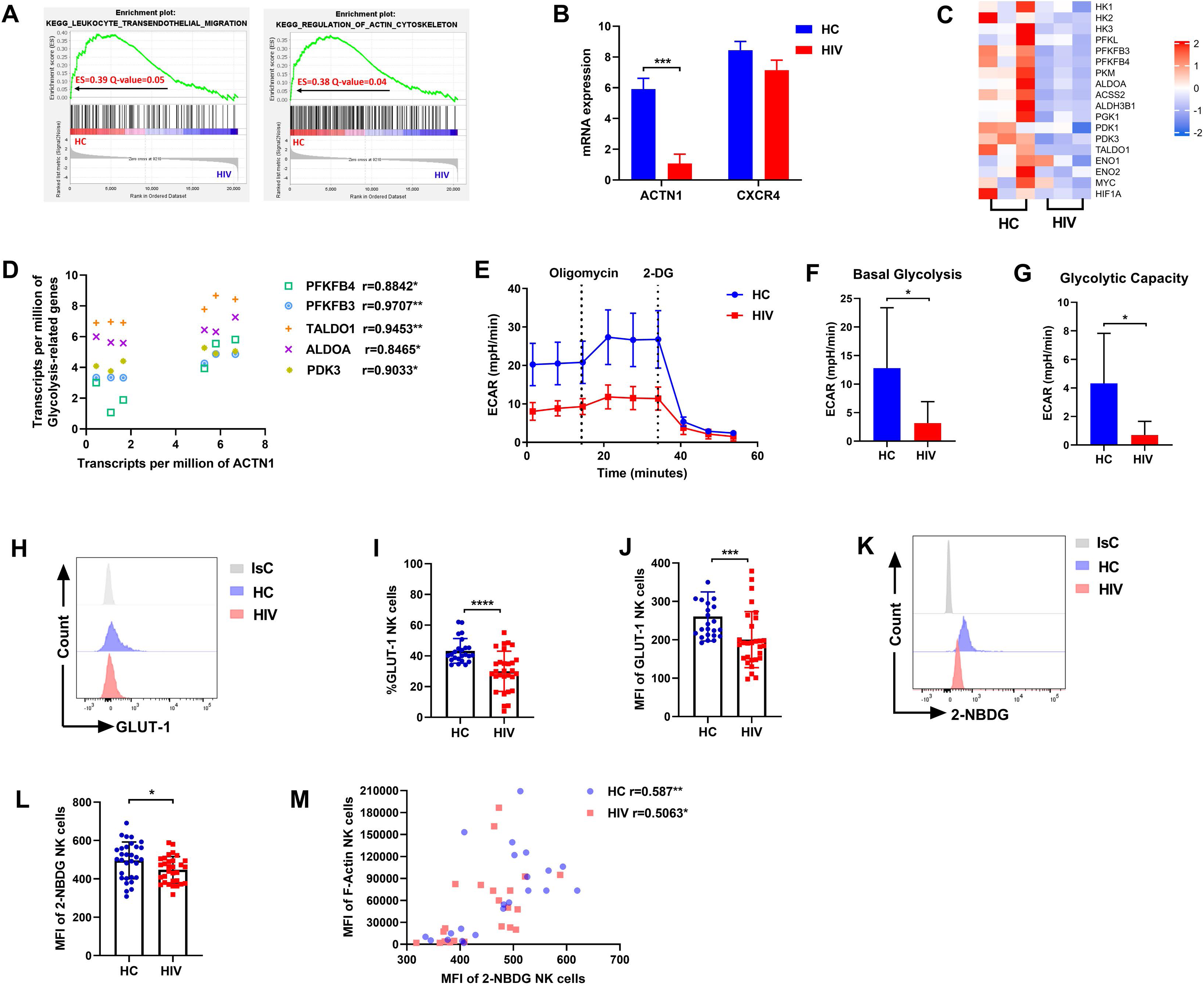
NK cell glycolysis is impaired in HIV infection. **(A-D)** Data from NCBI Gene Expression Omnibus (GEO) database identifier GSE25669. **(A)** GSEA analysis for KEGG lymphocyte trans-endothelial migration and regulation of Actin cytoskeleton in NK cells from HC and HIV individuals. Positive enrichment score (ES) indicates enrichment in HC individuals. Data from NCBI Gene Expression Omnibus (GEO) database identifier GSE25669. **(B)** Comparison of gene expression in HC (n = 3) and HIV (n = 3) individuals. **(C)** Heat map shows expression of glycolysis-associated gene expression of NK cells in HIV-infected individuals versus those in HC individuals. **(D)** The correlation between glycolysis-associated gene expression and *ACTN1* gene expression of NK cells (n = 6). **(E)** ECAR of NK cells in HC and HIV individuals. **(F and G)** The basal glycolysis (F) and the glycolytic capacity (G) of NK cells in HC (n = 6) and HIV (n = 6) individuals. **(H - J)** The percentage and MFI of GLUT-1 on NK cells from the HC (n = 24) and HIV (n = 29) individuals. **(K and L)** The MFI of 2-NBDG in NK cells from the HC (n = 31) and HIV (n = 31) individuals. **(M)** The correlation between 2-NBDG and F-Actin of NK cells from the HC (n = 22) and HIV (n = 22) individuals.

Subsequently, we found that the levels of glycolysis-related genes in NK cells of HIV-infected individuals were lower than those of HC individuals (Fig 2C), including hexokinase 1 (*HK1*), hexokinase 2 *(HK2*), 6-phosphofructo-2-kinase/fructose -2,6-bisphosphatase 4 (*PFKFB4*), 6-phosphofructo-2-kinase/fructose-2,6-bisphosphatase 3 (*PFKFB3*), Transaldolase (*TALDO1*), Fructose-bisphosphate aldolase A (*ALDOA*), phosphoglycerate kinase 1 (*PGK1*), pyruvate dehydrogenase kinase 1 (*PDK1*), and pyruvate dehydrogenase kinase 3 (*PDK3*). Moreover, we found that the transcripts per million of five glycolysis-related genes were significantly positively correlated with *ACTN1* by correlation analysis, including *PFKFB4*,*PFKFB3*,*TALDO1*,*ALDOA* and *PDK3* (Fig 2D). *ACTN1* encodes alpha-actin-1, a member of the actin crosslinked protein superfamily, which affects cell adhesion and movement by regulating the actin cytoskeleton^30–32^. Therefore, we speculated that impaired NK cell migration might be related to glycolysis in HIV infection, we examined the changes in glycolysis of NK cells during HIV infection.

To further examine differences in NK cell glycolysis between HC and HIV-infected individuals, we performed a detailed analysis of the extracellular acidification rate (ECAR) level of NK cells, which reflects the overall glycolytic flux by detecting changes in proton concentration in the media surrounding cells in real time via the use of the Seahorse analyzer. The results showed that the extracellular acidification rate (Fig 2E), basal glycolysis (Fig 2F) and glycolytic capacity (Fig 2G) of NK cells in HIV-infected individuals were significantly lower than those in HC individuals. NK cells need to ingest glucose to participate in glycolysis^28^, so we detected the expression of glucose transporter GLUT-1 on the surface of NK cells in HIV-infected individuals and HC individuals. The results found that the percentage and MFI of GLUT-1 expression on NK cells in HIV-infected individuals were significantly lower than those in HC individuals (Figs 2H, 2I and 2J). Furthermore, we assessed the glucose uptake capacity of NK cells using the fluorescent glucose analog 2-NBDG, which enters the cell through glucose transporters and is subsequently phosphorylated by hexokinase and accumulates in the cytoplasm as 2-NBDG-6-phosphate^33, 34^. The results showed that 2-NBDG uptake by NK cells in HIV-infected individuals was lower than that in HC individuals (Figs 2K and 2L). To explore whether the cytoskeletal movement of NK cells is related to glycolysis, we analyzed the correlation between F-Actin and 2-NBDG. We found that F-Actin levels in NK cells of HC individuals and HIV-infected individuals were significantly positively correlated with 2-NBDG uptake (Fig 2M), indicating that glycolysis might affect the movement of NK cells.

To investigate whether NK cell migration is controlled by the modulation of glycolysis, we compared NK cell migration in glucose-containing and glucose-depleted medium. As shown in Fig 3A, glucose depletion limits the migration of NK cells. Moreover, we assessed NK cell migration exposed to glycolysis inhibitor 2-DG and found that was significantly inhibited, especially 2-DG downregulated NK cell migration in a concentration-dependent manner (Figs 3B and 3C). In addition, we found that CXCL12-induced F-Actin elevation was prevented by 2-DG (Figs 3D and 3E). In summary, these results suggest that the impaired migration of NK cells in HIV-infected individuals is caused by the decrease of NK cell glycolysis.

**Fig 3.**
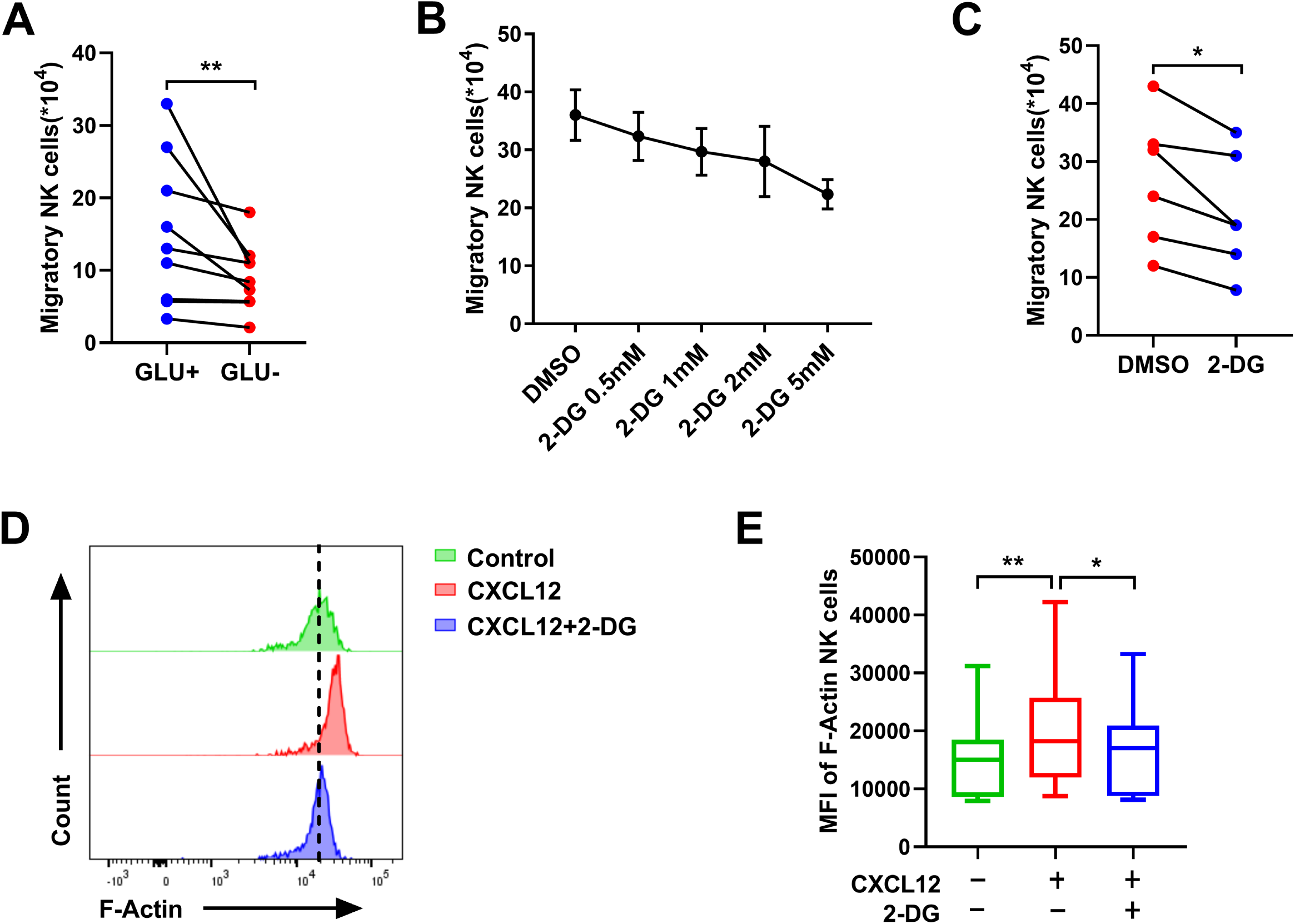
Engagement of glycolysis is necessary for NK cell migration. **(A)** CXCL12-triggered migration of NK cells in either glucose-containing or glucose-depleted medium (n = 9). **(B)** CXCL12-triggered migration of NK cells treated with different doses of 2-DG (0.5, 1, 2 and 5 mM) or DMSO (n = 3). **(C)** CXCL12-triggered migration of NK cells treated with 2-DG (2 mM) or DMSO (n = 6). **(D)** A representative flow cytometry plot demonstrating the effects of 2-DG (2 mM) or DMSO on the levels of F-Actin in CXCL12-stimulated NK cells. **(E)** The MFI of F-Actin in CXCL12-stimulated NK cells treated with 2-DG (2 mM) or DMSO (n = 9).

### PI3K/AKT/mTORC1 or ERK signaling pathway is involved in NK cell migration

Based on the above results, we intend to explore the metabolic pathways which link the glycolysis that happens during NK cell migration to the changes of actin. It has been reported that CXCL12 can induce the migration of endothelial cells and bone marrow mesenchymal stem cells via activating the PI3K/AKT signaling pathway^35, 36^. Studies have confirmed that the activation of AKT/mTORC1 signaling pathway can promote the glycolysis and effector functions of NK cells^28, 37^. Consequently, we speculate that the glycolysis-related signaling pathway PI3K/AKT/mTORC1 may be involved in NK cell migration. To verify this conjecture, we detected the phosphorylation levels of AKT and mTORC1 proteins in CXCL12-treated NK cells, and confirmed that there was an increase in phosphorylated AKT and phosphorylated S6 ribosomal protein (pS6), a readout of mTORC1 signaling (Figs 4A and 4B). In addition, we also demonstrated that CXCL12 could up-regulate the phosphorylation level of ERK in NK cells (Figs 4C and 4D).To address if glycolysis-related PI3K/AKT/mTORC1 or ERK pathway underpins the migration of NK cells, we investigated the levels of F-Actin in cultured NK cells treated with inhibitors of the PI3K/AKT/mTORC1 or ERK signaling pathway (Fig 4E) and found that inhibitors of AKT, mTORC1 and ERK all can significantly inhibit the level of F-Actin in NK cells (Figs 4F and 4G). The above results indicate that CXCL12 can promote the migration of NK cells by activating the glycolysis-related PI3K/AKT/mTORC1 or ERK signaling pathway.

**Fig 4.**
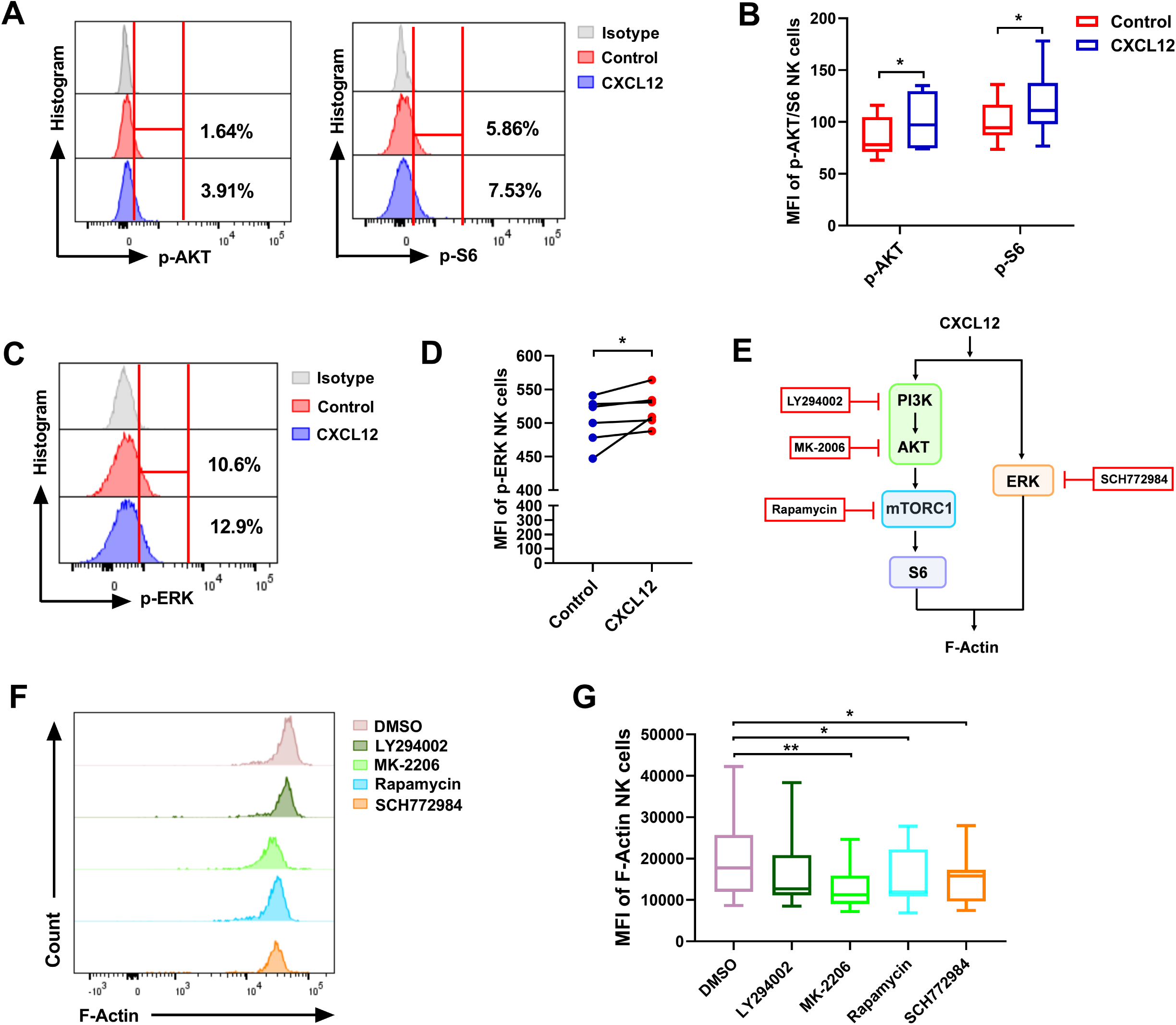
PI3K/AKT/mTORC1 or ERK signaling pathway is involved in NK cell migration. **(A)** The representative flow cytometry plots show the effects of CXCL12 (100 ng/ml) on the phosphorylation of AKT and S6 in NK cells. **(B)** The phosphorylation MFI of AKT and S6 in NK cells stimulated with CXCL12 (100 ng/ml) (n = 6). **(C)** The representative flow cytometry plots show the effects of CXCL12 (100 ng/ml) on the phosphorylation of ERK in NK cells. **(D)** The phosphorylation MFI of ERK in NK cells stimulated with CXCL12 (100 ng/ml) (n = 6). **(E)** Schematic diagram of inhibitors in PI3K/AKT/mTORC1 and ERK signaling pathways. **(F)** A representative flow cytometry plot demonstrating the effects of LY294002 (50 μM), MK-2206 (10 μM), Rapamycin (200 nM), SCH772984 (300 nM) or DMSO on the levels of F-Actin in CXCL12-stimulated NK cells. **(G)** The MFI of F-Actin in CXCL12-stimulated NK cells treated as indicated (n = 9).

### High expression of TIGIT in HIV-infected individuals inhibits glycolysis of NK cells

In view of the fact that NK cell migration is regulated by glycolysis and glycolysis-related signaling pathways, we then explore the possible reasons for the decline of NK cell glycolysis in HIV-infected individuals. It has been found that T cell metabolism is regulated by inhibitory receptors in recent years^38, 39^, so it needs to investigate whether the impaired glycolysis of NK cells in HIV infection is related to inhibitory receptors. We detected the expression of three common inhibitory receptors on the surface of NK cells by flow cytometry. The results showed that the expression of T cell immunoglobulin and ITIM domain (TIGIT) on NK cells of HIV-infected individuals was higher than that of HC individuals, while there was no significant difference in the expression of lymphocyte-activation gene 3 (LAG-3) and cytotoxic T-lymphocyte-associated protein 4 (CTLA-4) on NK cells between these two groups (Figs 5A and 5B). Therefore, we investigated the effect of TIGIT expression on NK cell glycolysis in HC and HIV-infected individuals. The results showed that the expression of GLUT-1 was significantly lower in TIGIT^+^ NK than TIGIT^-^ NK cells (Figs 5C and 5D). Additionally, 2-NBDG was negatively correlated with TIGIT expression in NK cells (Fig 5E). We next investigated whether TIGIT blockade could restore the decrease in NK cell glycolysis during HIV infection, and found that the levels of GLUT-1 (Figs 5F and 5G), 2-NBDG (Figs 5H and 5I), and ECAR (Fig 5J) were increased by blocking TIGIT.

**Fig 5.**
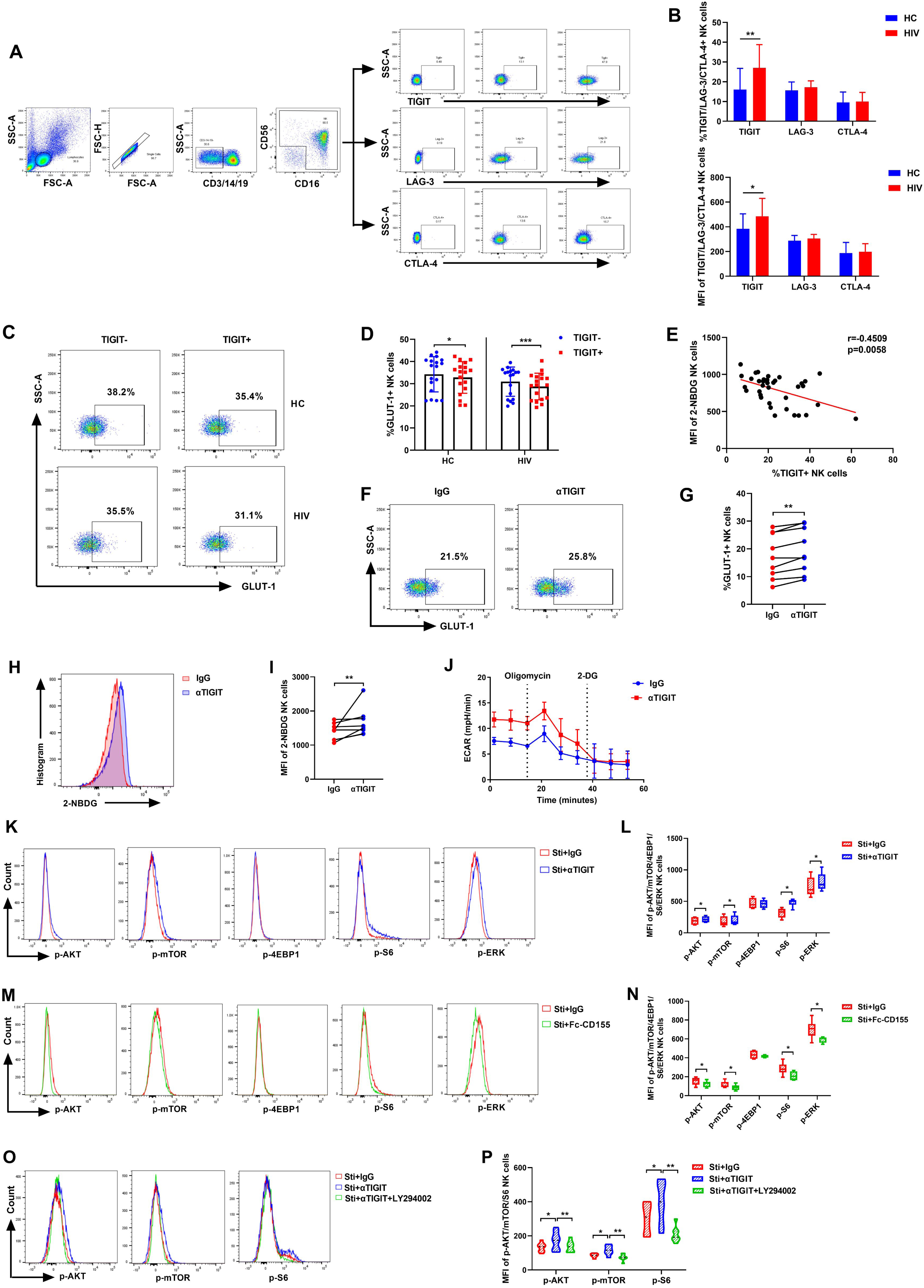
Increased TIGIT-positive NK cells disrupt the glycolysis of HIV-infected individuals by inhibiting PI3K/AKT/mTORC1 or ERK signaling pathways. **(A)** The representative flow cytometry plots showing the expression of TIGIT, LAG-3 and CTLA-4 on NK cells from HC and HIV individuals. **(B)** The percentage and MFI of TIGIT, LAG-3 and CTLA-4 on NK cells from HC (n = 18) and HIV (n = 18) individuals. **(C and D)** The percentages of GLUT-1on TIGIT ^-^ or TIGIT ^+^ NK cells from HC (n = 18) and HIV (n = 18) individuals. **(E)** The correlation between TIGIT and 2-NBDG of NK cells (n = 36). **(F)** The representative flow cytometry plots show the effects of αTIGIT (5 μg/ml) on the expression of GLUT-1 on NK cells. **(G)** The percentage of GLUT-1 on NK cells treated with αTIGIT (5 μg/ml) or isotype control (n = 9). **(H)** The representative flow cytometry plots show the effects of αTIGIT (5 μg/ml) on the glucose uptake in NK cells. **(I)** The MFI of 2-NBDG in NK cells treated with αTIGIT (5 μg/ml) or isotype control (n = 8). **(J)** ECAR of NK cells treated with αTIGIT (5 μg/ml) or isotype control. **(K and M)** The representative flow cytometry plots show the effects of αTIGIT (5 μg/ml) (K) and Fc-CD155 (5 μg/ml) (M) on the phosphorylation of AKT, mTOR, 4EBP1, S6 and ERK in NK cells. **(L and N)** The phosphorylation MFI of AKT, mTOR, 4EBP1, S6 and ERK in NK cells treated with αTIGIT (5 μg/ml) (L) and Fc-CD155 (5 μg/ml) (N) (n = 6). **(O)** The representative flow cytometry plots show the effects of LY294002 (50 μM) on the phosphorylation of AKT, mTOR and S6 in αTIGIT-treated NK cells. **(P)** The phosphorylation MFI of AKT (n = 8), mTOR (n = 7) and S6 (n = 7) in αTIGIT-treated NK cells treated with LY294002 (50 μM).

As data shown above that TIGIT can inhibit NK cell glycolysis, we continue to explore how TIGIT affect NK cell glycolysis. Our results above showed that NK cell migration is regulated by the PI3K/AKT/mTORC1 or ERK signaling pathways, therefore, we studied the effect of TIGIT on the activity of AKT/mTORC1 and ERK signaling pathways. As expected, AKT, mTOR, S6 and ERK phosphorylation were increased in NK cells after TIGIT blockade compared to isotype control (Figs 5K and 5L). We treated NK cells with exogenous CD155, a ligand of TIGIT, and found that AKT, mTOR, S6 and ERK phosphorylation were down-regulated (Figs 5M and 5N). Moreover, inhibition of PI3K activity decreased the phosphorylation of AKT, mTOR and S6, although TIGIT is blocked (Figs 5O and 5P). These results suggest that TIGIT inhibits NK cells glycolysis through PI3K/AKT/mTORC1 or ERK signaling pathways.

### TIGIT suppresses NK cell migration by inhibiting glycolysis through HIF-1α pathway

Given that TIGIT could weaken the glycolysis of NK cells, and inhibition of glycolysis could damage the migration of NK cells, we thus explore the role of TIGIT on NK cell migration next. We evaluated the expression of F-Actin involved in migration process by Flow cytometry first. Expression of F-Actin was significantly reduced in TIGIT ^+^ NK cells compared with TIGIT ^-^ NK cells (Figs 6A and 6B), and we found that the levels of F-Actin expression and migration of NK cells were increased by blocking TIGIT (Figs 6C-6E). Also, we observed that the trigger of exogenous CD155 weakened NK cell migration (Fig 6F).

**Fig 6.**
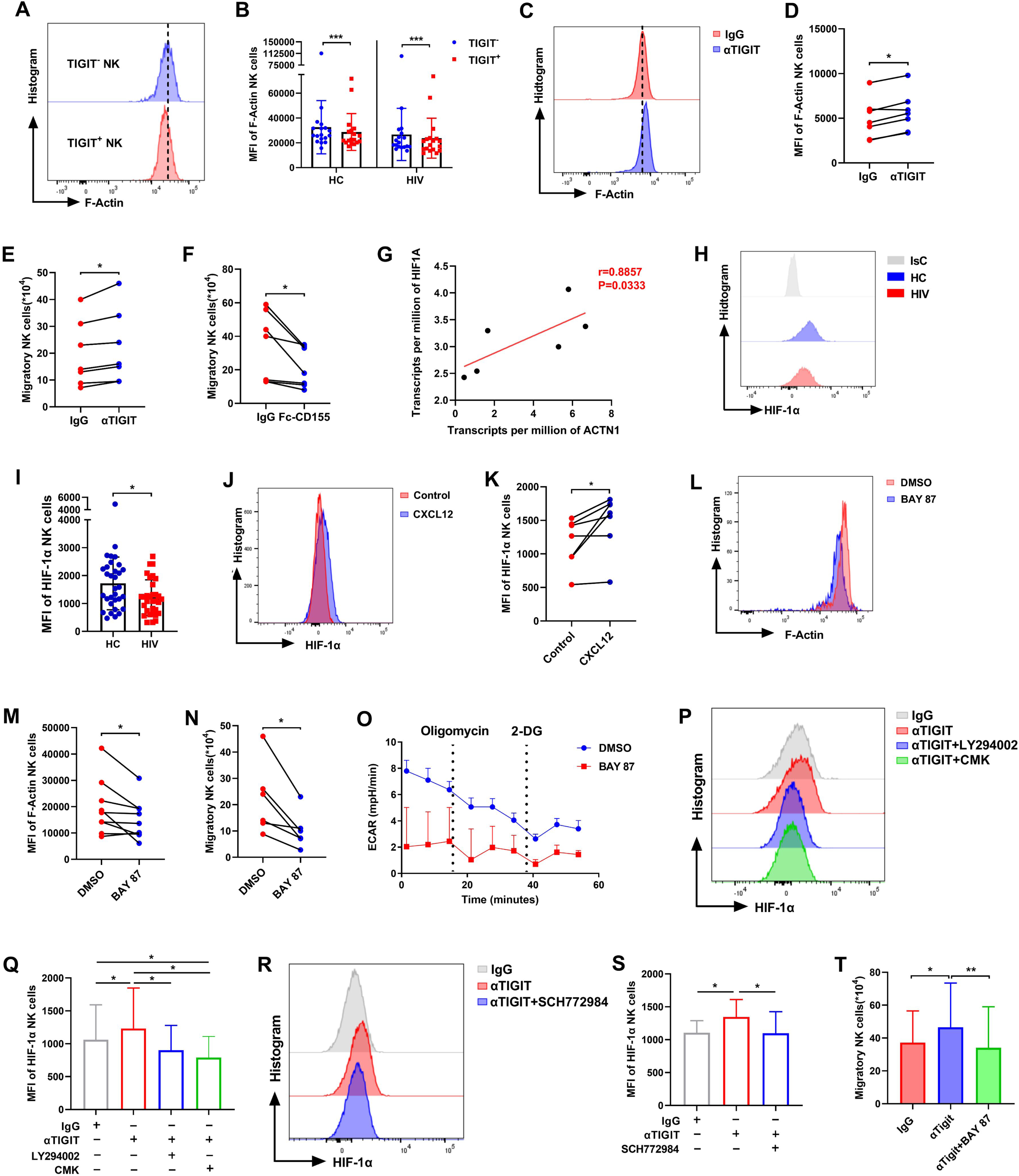
TIGIT suppresses NK cell migration via inhibiting HIF-1α-mediated glycolysis. **(A and B)** The MFI of F-Actin in TIGIT^-^ or TIGIT^+^ NK cells from HC (n = 18) and HIV (n = 18) individuals. **(C)** A representative flow cytometry plot demonstrating the effects of αTIGIT (5 μg/ml) on the levels of F-Actin in CXCL12-stimulated NK cells. **(D)** The MFI of F-Actin in CXCL12-stimulated NK cells treated with αTIGIT (5 μg/ml) or isotype control (n = 7). **(E and F)** CXCL12-triggered migration of NK cells treated with αTIGIT (5 μg/ml) (E) and Fc-CD155 (5 μg/ml) (F) (n = 7). **(G)** The correlation between *HIF1A* gene expression and *ACTN1* gene expression of NK cells (n = 6). Data from NCBI Gene Expression Omnibus (GEO) database identifier GSE25669. **(H and I)** The MFI of HIF-1α in NK cells from the HC (n = 32) and HIV (n = 32) individuals. **(J)** A representative flow cytometry plot shows the effects of CXCL12 (100 ng/ml) on the expression of HIF-1α in NK cells. **(K)** The MFI of HIF-1α in NK cells stimulated with CXCL12 (100 ng/ml) (n = 7). **(L)** A representative flow cytometry plot demonstrating the effects of BAY 87 (10 μM) or DMSO on the levels of F-Actin in CXCL12-stimulated NK cells. **(M)** The MFI of F-Actin in CXCL12-stimulated NK cells treated with BAY 87 (10 μM) or DMSO (n = 9). **(N)** CXCL12-triggered migration of NK cells treated with BAY 87 (10 μM) or DMSO (n = 6). **(O)** ECAR of NK cells treated with BAY 87 (10 μM) or DMSO. **(P)** A representative flow cytometry plot shows the effects of LT294002 (50 μM) and CMK (50 μM) on the expression of HIF-1α in αTIGIT-treated NK cells. **(Q)** The MFI of HIF-1α in αTIGIT-treated NK cells treated with LT294002 (50 μM), CMK (50 μM) or DMSO (n = 6). **(R)** A representative flow cytometry plot shows the effects of SCH772984 (300 nM) on the expression of HIF-1α in αTIGIT-treated NK cells. **(S)** The MFI of HIF-1α in αTIGIT-treated NK cells treated with SCH772984 (300 nM) or DMSO (n = 7). **(T)** CXCL12-triggered migration of NK cells treated with αTIGIT (5 μg/ml) and BAY 87 (10 μM) (n = 7).

We further delve deeper into the mechanism by which TIGIT inhibits NK cell migration. By correlation analysis at the gene level, we found that the mRNA level of *HIF1A* was significantly positively correlated with *ACTN1* (Fig 6G). We detected the level of Hypoxia-inducible factor-1 alpha (HIF-1α) by flow cytometry, and the result showed that the protein expression level of HIF-1α in NK cells of HIV-infected individuals was lower than that of HC individuals (Figs 6H and 6I). Next, we demonstrated increased HIF-1α expression in NK cells after the treatment of CXCL12 (Figs 6J and 6K), and found that HIF-1α inhibitor BAY 87 significantly inhibited F-Actin level (Figs 6L and 6M) and NK cell migration (Fig 6N). These findings suggested that the HIF-1α pathway is involved in NK cell migration.

Moreover, by detecting ECAR, it was found that the glycolysis level of NK cells dramatically declined by inhibiting HIF-1α (Fig 6O). And the expression of HIF-1α in NK cells increased by blocking TIGIT compared to isotype control (Figs 6P-6S). Inhibitors for PI3K/S6K or ERK signaling pathway decreased the expression of HIF-1α in NK cells although TIGIT is blocked (Figs 6P-6S). These results displayed that TIGIT inhibits NK cells glycolysis through PI3K/AKT/mTORC1/HIF-1α or ERK/HIF-1α signaling pathways.

Finally, we explored the link between HIF-1α-mediated glycolysis and motility affected by TIGIT by analyzing the migration of NK cells. And we found that inhibition of HIF-1α activity decreased the migration of NK cells although TIGIT is blocked (Fig 6T), implying that TIGIT suppresses NK cell migration via inhibiting HIF-1α-mediated glycolysis.

## Discussion

NK cells make an important contribution to the resistance to HIV infection, and the function of NK cells is closely related to their migration ability. However, the migration of NK cells and the relationship between NK cell migration and disease progression in HIV-infected individuals are unclear. It was found in the present study that the migration of NK cells in HIV-infected individuals is significantly weaker than that in HC individuals. Additionally, we found that the decreased migration of NK cells is associated with the increased severity of disease. It was reported that the immunosuppressive microenvironment in the tumor weakens the function of NK cells and the movement of NK cells^40^. And in a variety of solid tumors, the NK cell infiltration can be used as a prognostic factor in the survival of tumor patients^41–44^. Salazar-Mather et al. reported that NK cells can exert optimal anti-viral defense when recruited to the inflammatory sites during murine cytomegalovirus (MCMV) infection^22^. Therefore, the decreased NK cell migration may result in reduced contact between NK cells and infected cells, thus weakening the role of NK cells in resisting viral infection, and attenuating the therapeutic outcomes of HIV-infected individuals.

The present study found that it is the reduction of glycolysis of NK cells in HIV-infected individuals that weakens the migration of NK cells. Recently research has highlighted the importance of glycolysis governing the cytokine secretion and killing effect of NK cells^24, 27, 45–47^. Our results showed that glycolysis also plays an important role in regulating NK cell migration. Several previous studies have also shown that the migration of T cells and dendritic cells requires the participation of glycolysis^48–50^, which provides data and theoretical support for our conclusion. The glycolysis of NK cells takes glucose as fuel, and glucose-driven glycolysis promotes the anti-viral and anti-tumor effects of NK cells^45, 46^. We found that glucose depletion significantly inhibited NK cell motility. Kishore et al. found a similar situation in Treg cells, where glucose depletion also inhibited Treg cell migration^49^. Moreover, the cytoskeletal component actin can interact with glycolytic enzymes^51^ and form a glycolytic ATP donor for the ATP-hydrolyzing sodium pump, thus providing energy for cytoskeletal rearrangement^52^. Kishore et al. also demonstrated that glucokinase (GCK) promotes cytoskeletal rearrangement and migration of Treg cells by associating with actin^49^. Thus, we hypothesized that the possible reason for the restriction of cytoskeleton rearrangement and migration of NK cells caused by glycolytic inhibition due to the suppression of the interaction between glycolytic enzymes with actin. In addition, Haas et al. reported that chemokines up-regulate the expression of hexokinase HK1 and pyruvate kinase PKM1/2, the glycolysis rate-limiting enzymes, at both transcriptional and translational levels during the process of CD4^+^ T cell migration, and that exposure of CD4^+^ T cells to chemokines resulted in increased glucose transporter gene level and glycolysis flux^48^. Therefore, we speculated that chemokines acting on NK cells may also induce glycolysis of NK cells by up-regulating key glycolytic enzymes or glycolysis-related signaling pathways, thus promoting NK cell migration.

We also observed the glycolysis-related signaling pathway PI3K/AKT/mTORC1 or ERK is involved in NK cell migration. In a variety of cells, the signaling pathway controlled by PI3K/AKT regulates cell metabolism, cell growth, proliferation, actin cytoskeleton rearrangement and migration^53–55^. mTOR-mediated signaling pathway is one of the key downstream signaling pathways regulated by AKT, which controls cell response to environmental stress stimuli through the formation of two different complexes, mTORC1 and mTORC2^56^. Kishore et al. found that pro-migration stimulators initiate the glycolysis of Treg cells by activating the PI3K/AKT/ mTORC2-mediated signaling pathway, thereby promoting cytoskeletal rearrangement and migration^49^. Moreover, PI3K/AKT/mTOR and ERK signaling pathways have been reported as common downstream signaling pathways with CXCR4 activation^57–60^. Although the metabolic pathways involved in the migration of human NK cells have not been elucidated previously, in mice, Saudemont et al. demonstrated that two subtypes of PI3K, p110γ and p110δ, are essential for the chemotaxis of mouse NK cells to CXCL12^61^, which also supported our conclusion.

We further explored the causes of impaired glycolysis and found that high expression of TIGIT in NK cells inhibited glycolysis of HIV-infected individuals. It is well known that the imbalance between inhibitory receptors and activated receptors of HIV-infected individuals will lead to immune cell dysfunction^11, 62^. In recent years, inhibitory receptors have been shown to be potentially useful targets in HIV immunotherapy. There are differences in the expression of TIGIT on normal human peripheral blood immune cells, and Wang et al. have demonstrated that TIGIT expression on NK cells is relatively high, so it may be more important in regulating the effector function of NK cells^63^. Our study showed that TIGIT expression on NK cells of HIV-infected individuals was significantly increased, which was consistent with the previous research results of our team, and the previous research of our team also found that the high expression of TIGIT on NK cells could inhibit the production of IFN-γ in HIV-infected individuals^64^. Moreover, during HIV infection, the cytokine secretion function of TIGIT^+^ NK cells was weaker than that of TIGIT^-^ NK cells^65^, and blocking TIGIT could restore the cytotoxicity of NK cells^66^. However, whether TIGIT affects the metabolism of NK cells has not attracted much attention, our study revealed for the first time that TIGIT played an inhibitory effect on NK cell glycolysis. Furthermore, we discovered that TIGIT restricts the activation of glycolysis-related pathways in NK cells and reduces the phosphorylation of PI3K/AKT/mTORC1 and ERK signaling pathways. This is possibly because the binding of TIGIT to CD155 promotes TIGIT cytoplasmic tail phosphorylation through Src family kinases Fyn and Lck, leading to recruitment of SHIP-1 (inositol 5-phosphatase-1 containing the SH2 domain), thus inhibiting downstream PI3K/AKT and ERK signaling pathways^67–70^, resulting in weakened NK cell glycolysis. Similar to our study, He et al. showed that TIGIT reduces glucose uptake of CD8^+^ T cells by inhibiting the activity of the AKT/mTOR signaling pathway, thereby damaging the cell metabolism and effector function of CD8^+^ T cells in Gastric Cancer^38^.

Furthermore, we discovered that TIGIT inhibits the cytoskeletal rearrangement and migration of NK cells by impairing glycolysis, remarkably, HIF-1α is a key molecule in this process. HIF-1α induces the gene expression of glucose transporters and glycolytic enzymes to initiate glycolysis^15, 71^. Therefore, we speculated that HIF-1α participates in glycolysis of NK cells perhaps by regulating glucose uptake or the expression of glycolytic enzymes. Moreover, we provided evidence that TIGIT inhibits HIF-1α expression in NK cells through the PI3K/AKT/mTORC1 or ERK signaling pathway. Previous studies have shown that PI3K and AKT signals are involved in promoting HIF-1α synthesis in prostate cancer cells^72^, and mTORC1 increases glycolytic flux by activating HIF-1α transcription and translation^73, 74^. In addition, blocking the ERK/HIF-1α/GLUT-1 pathway was found to down-regulate macrophage glycolysis in a rat rheumatoid arthritis model^75^. Our study also reported that HIF-1α-mediated glycolysis promotes the migration of NK cells. Liu et al. found that chemokines activate the HIF-1α signaling pathway in dendritic cells, leading to metabolic reprogramming towards glycolysis, thereby promoting dendritic cell migration^50^. Finlay et al. demonstrated that mTORC1/HIF-1 pathway is necessary for the maintenance of glycolysis in effective cytotoxic T lymphocytes (CTL), and this pathway regulates the expression of essential chemokines and adhesion receptors that mediate CD8^+^ T cell transport^76^. These findings provide strong support for our conclusions. We also found that blocking TIGIT restored the impaired glycolysis and migration of NK cells in HIV-infected individuals. Combined with previous research results, blocking TIGIT can restore the immune response of NK cells and the cytotoxic activity of NK cells against the HIV reservoir^65, 66, 77^, highlighting the potential of TIGIT blockade in immunotherapy HIV treatment strategies.

In conclusion, our results highlight the suppression of glycolysis inhibited the migration of NK cells during HIV infection. This mechanism may be controlled by the decreased of HIF-1α expression mediated by PI3K/Akt/mTORC1 or ERK pathways regulated via TIGIT. (Fig. 7). As far as we know, the present study is the first to investigate the mechanism by which migration and glycolysis of NK cells are inhibited in HIV infection. This discovery could provide a scientific theoretical basis for the immunotherapy of HIV-infected individuals and promote to identify therapeutic targets for the improvement of immune function in HIV infection.

**Fig 7.**
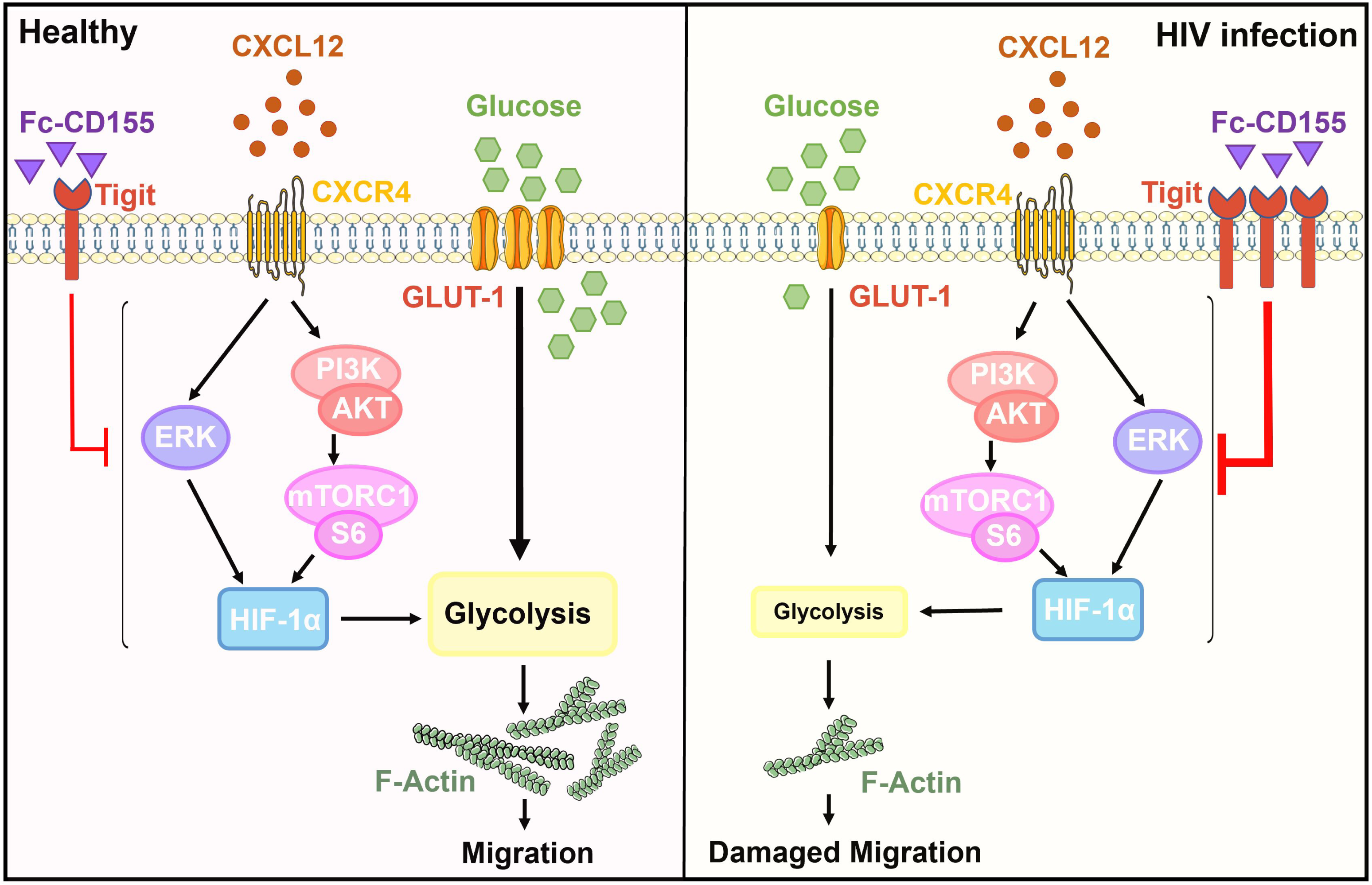
Schematic of the mechanism of regulating NK cell migration in HC and HIV-infected individuals. Under normal physiological conditions, NK cells express enough glucose transport receptor GLUT-1 to support glycolysis, and the glucose uptake capacity is normal. Meanwhile, the activation of PI3K/AKT/mTORC1 or ERK signaling pathways can induce HIF-1α expression, subsequently promotes intracellular glycolysis and maintains the normal movement of NK cells. During HIV infection, the expression of GLUT-1 on NK cells is down-regulated, resulting in abnormal glucose uptake. Highly expressed TIGIT limits HIF-1α expression by inhibiting the PI3K/AKT/ mTORC1 or ERK signaling pathway, leading to impaired glycolysis, thereby suppressing the migration of NK cells.

## Materials and methods

### Study participants

Peripheral blood was collected from 168 HIV-infected individuals from State Key Laboratory for Diagnosis and Treatment of Infectious Diseases, NHC Key Laboratory of AIDS Prevention and Treatment, The First Affiliated Hospital of China Medical University. HIV-infected individuals included ART-naïve HIV-infected individuals, immunological responders (IRs), and Immunological non-responders (INRs). IRs were HIV-infected individuals on antiretroviral therapy with CD4^+^ T cell counts greater than 500 cells/μl; INRs were those with CD4^+^ T cell counts less than 350 cells/μl. 152 Healthy Control (HC) individuals (matched for age and sex with HIV-infected individuals) were enrolled in this study. The inclusion criteria of HC individuals were exclusion of infection and autoimmune diseases. The study was approved by the Research and Ethics Committee of The First Affiliated Hospital, China Medical University.

### Cell isolation and cell culture

Blood samples were collected from HIV-infected individuals or age- and sex-matched Healthy Control (HC) individuals. Peripheral blood mononuclear cells (PBMCs) extracted from fresh blood were isolated by Ficoll-Hypaque density gradient centrifugation. Total NK cells were purified from PBMCs by negative selection using the EasySep™ Human NK Cell Enrichment Kit (STEMCELL, Canada). Purified NK cells were cultured in RPMI 1640 medium supplemented with 10% heat-inactivated fetal bovine serum (FBS), penicillin (50 U/ml) (Invitrogen, USA), and streptomycin (50 μg/ml) (Invitrogen, USA) in the incubator at 37°C with 5% CO2.

### Transwell Migration Assay

The migration assay was performed using 24-well Transwell plates containing 8 μm pore size polycarbonate filter (Corning, NY). The chemokine CXCL12 binds to CXCR4, which is highly expressed by NK cells, to induce NK cell migration^78^. Migration medium or chemokines CXCL12 (100 ng/ml; R&D, USA) were added to the lower chambers. NK cells were added to the upper chamber and incubated at 37°C for 3 h. Then the upper chamber was removed and NK cells in the lower chamber were counted by hemocytometer. In some experiments, NK cells were pretreated with DMSO or 2-DG (0.5 mM, 1 mM, 2 mM, 5 mM; Sigma-Aldrich, USA), BAY 87 (10 μM; Cayman, Germany) for 3 h. In other experiments, NK cells were pretreated with Recombinant Human CD155 (5 μg/ml; R&D, USA), anti-TIGIT blocking antibody (5 μg/ml; eBioscience, USA), or IgG control for 1h.

### Detection of Absolute CD4^+^ T Cell Counts

The TriTEST anti-CD4-FITC/CD8-PE/CD3-PerCP reagents (BD Biosciences, USA), anticoagulated whole blood and 1×disposable hemolysin were added to the Trucount tubes for a single-platform lyse-no-wash procedure. The cell counts were detected by flow cytometer (BD FACS Calibur), and the results were analyzed by MultiSET software.

### Measurement of F-Actin and phosphorylated cofilin

NK cells were resuspended in BD Cytofix/Cytoperm Solution (BD Biosciences, USA) for 30 min at 4℃ in the dark, then washed 2 times with 1× BD Perm/Wash™ buffer (BD Biosciences, USA). F-Actin was labeled with Alexa Fluor 647 Phalloidin (Thermo Fisher Scientific, USA) for 30 min at 4℃ in the dark. Then the cells were washed with BD Perm/Wash Buffer and detected by flow cytometry. In some experiments, NK cells were prepared overnight with a number of drugs purchased from Sigma-Aldrich, MCE or Cayman: 2-DG (5 mM), Rapamycin (200 nM), LY294002 (50 μM), MK-2206 (10 μM), SCH772984 (300 nM), BAY 87 (10 μM). Then the cells were stimulated with CXCL12 (100 ng/ml) at 37°C for 15 min. After treatment of NK cells with CXCL12, cells were washed with cold phosphate-buffered saline (PBS) with 2% fetal bovine serum and then stained by flow cytometry. BD Horizon™ Fixable Viability Stain 620 (BD Biosciences, USA) was used to evaluate cell viability.

Cofilin dynamics were measured as the percentage and mean fluorescence intensity (MFI) of phospho-cofilin. NK cells were fixed with 1.5% formaldehyde and incubated at room temperature for 10min. Then the cells were permeabilized with cold (−20℃) methanol on ice for 10min. The cells were next stained with rabbit anti-p-cofilin (Ser3) antibody (Cell Signaling Technology, USA) for 1 h and washed 2 times. The cells were labeled with Alexa Fluor 488-conjugated chicken anti-rabbit antibody (Invitrogen, USA) for 30 min and then detected by flow cytometry.

### Cell surface and intracellular staining

The phenotype of NK cells was analyzed from freshly isolated PBMCs by flow cytometry. CXCR4 and GLUT-1 expression on NK cells were detected using the following fluorophore-conjugated antibodies: FITC anti-human CD3, Percp-Cy5.5 anti-human CD14, Percp-Cy5.5 anti-human CD19, PE-Cy7 anti-human CD56, APC-Cy7 anti-human CD16, BV421 anti-human CXCR4, BV421 Mouse IgG2a κ isotype control (all from BD biosciences, USA), PE anti-human GLUT-1(R&D, USA), and PE Mouse IgG2B isotype control (R&D, USA). TIGIT, LAG-3, and CTLA-4 expression on NK cells were detected using the following fluorophore-conjugated antibodies: Percp-Cy5.5 anti-human CD3/CD14/CD19, PE-Cy7 anti-human CD56, APC-Cy7 anti-human CD16, APC anti-human CTLA-4, APC Mouse IgG2a κ isotype control (all from BD biosciences, USA), BV421 anti-human TIGIT, BV421 Mouse IgG2a κ isotype control, FITC anti-human LAG-3, and FITC Mouse IgG1 κ isotype control (all from Biolegend, USA). In brief, freshly isolated PBMCs were labeled with the corresponding fluorescently antibodies cocktails for 30 min at 4°C. Then stained cells were washed with PBS containing 2% FBS and detected by LSR II flow cytometer (BD Biosciences). The data was analyzed by FlowJo v10 software (Ashland, OR, USA).

For intracellular HIF-1α staining, eBioscience™ Foxp3 / Transcription Factor Staining Buffer Set was used (Thermo Fisher Scientific, USA). Cells were fixed/permeabilized for 45 min at 4°C with the prepared Fixation/Permeabilization working solution, which was prepared by mixing 1 part of fixation/permeabilization concentrate with 3 parts of fixation/permeabilization diluent. Cells were then washed 2 times with 1× permeabilization buffer and stained with PE anti-human HIF-1α antibody for 45 min at 4°C. The cells were finally washed with 1×permeabilization buffer, then centrifuged and resuspended in 200 μl PBS.

### Glucose uptake assay

2-NBDG (2-(N-(7-nitrobenzene-2-oxy-1, 3-diazole-4-yl) amino) -2-deoxyglucose) is a fluorescent glucose analogue used to monitor glucose uptake in cells. According to the manufacturer’s instructions, freshly isolated cells were resuspended in glucose-free R10 with 2-NBDG (Thermo Fisher Scientific, USA) at a final concentration of 100 μM for 30 minutes at 37℃. Finally, the cells were washed 2 times with warm PBS, and stained for surface markers at 4℃ for 30 minutes in a dark environment. The absorption of fluorescence by NK cells was detected immediately by LSR II flow cytometry.

### Extracellular acidification rate assays

A Seahorse XF HS Mini Analyzer (Agilent, USA) was used for real-time analysis of the extracellular acidification rate (ECAR) of purified NK cells cultured for 48 hours in 10 ng/ml IL-12, 50 ng/ml IL-15 and 100 ng/ml IL-18 (all from R&D, USA). In some experiments, NK cells were pretreated with anti-TIGIT blocking antibody (5 μg/ml; eBioscience, USA) or Isotype control for 1 h prior to stimulation. Briefly, 4×10^5^ treated NK cells were resuspended in Seahorse XF DMEM Medium containing glutamine (1 mM; Agilent, USA) and seeded into an 8-well XF cell culture microplate (Agilent, USA). All cell culture plates were treated with Cell-Tak (Corning, USA) for 20 minutes to ensure that NK cells adhered to the culture plates. Measurements of ECAR following addition of oligomycin (2 μg/ml; MCE, USA) and 2-deoxyglucose (50 mM; Agilent, USA) allowed for the calculation of basal glycolysis and glycolytic capacity. Each experimental sample was done in at least triplicate wells.

### Phosphorylation detection of signaling pathways

Freshly isolated NK cells were stimulated with or without chemokine CXCL12 (100 ng/ml; R&D, USA) for 15 min at 37℃ and 5% CO2. Alternatively, NK cells were stimulated with 10 ng/ml IL-12, 50 ng/ml IL-15 and 100 ng/ml IL-18 (all from R&D, USA) for 15 min after treatment with 5μg/ml anti-TIGIT blocking antibody (5 μg/ml; eBioscience, USA) or isotype control for 1 h. Then the cells were immediately fixed to stop stimulation with the same volume of warm BD Phosflow™ Fix Buffer I (BD biosciences, USA) for 10 min at 37℃. Cells were washed with PBS (2%FBS) and resuspended in 100ul ice-cold BD Phosflow™ Perm Buffer III (BD biosciences, USA) and incubated on ice for 30 min. Subsequently, the cells were washed twice with PBS (2%FBS) and stained for APC-S6 (eBioscience, USA), Alexa 647-AKT, PE-mTOR, Alexa 488-4EBP-1, and Alexa 488-ERK1/2 (all from BD biosciences, USA) for 30 min at 4 ℃. Finally, the cells were washed and resuspended in PBS (2%FBS) and immediately analyzed by LSR II flow cytometry.

### TIGIT Blockade and activation Assays

TIGIT was blocked by anti-TIGIT blocking antibody (αTIGIT, 5 μg/ml; eBioscience, USA) and activated by recombinant human CD155 (Fc-CD155, 5 μg/ml; R&D, USA). The effects of blocking and activating TIGIT on NK cells were assessed by pre-incubating cells in the presence of αTIGIT (5 μg/ml; eBioscience, USA), Fc-CD155 (5 μg/ml; R&D, USA) or isotype control for 1 h, followed by stimulation with 10 ng/ml IL-12, 50 ng/ml IL-15 and 100 ng/ml IL-18 (all from R&D, USA) for 15 minutes or 24 hours at 37◦C and 5% CO_2_. After incubation, signaling pathway phosphorylation, GLUT-1 expression and 2-NBDG uptake were evaluated as described above. For F-Actin detection, NK cells were cultured with αTIGIT antibody (5 μg/ml; eBioscience, USA) or isotype control for 1 h, and then stimulated with CXCL12 (100 ng/ml; R&D, USA) at 37◦C and 5% CO_2_ for 15 minutes.

### Online database analysis

We use the online resource GEO database (https://www.ncbi.nlm.nih.gov/geo/) to analyze the difference of NK cell RNA-sequencing (RNA-seq) data between HC individuals and HIV-infected individuals. The RNA-seq data set was from GEO with the number of GSE25669. TPM (transcripts per million) was used to measure gene expression levels. The data were normalized to TPM using the “limma” package in R v3.6.3 software (https://www.r-project.org/), and Log-2 transformations were performed on all data. GSEA was analyzed by GSEA v4.1.0 software (Broad Institute software). The enrichment pathways data set was analyzed using the Kyoto Encyclopedia of Genes and Genomes (KEGG) in the MSigDB data set. Heat map was plotted using phatmap and ComplexHeatmap package in R language.

### Statistical analysis

All flow cytometry data were analyzed by FlowJo software v10.6.2. Statistical analysis was performed using GraphPad Prism v8.0 software and SPSS Statistics v26.0 software programs. Two groups of data were analyzed by Wilcoxon matched-pairs signed rank tests or Mann-Whitney test. Kruskal-Wallis test or Friedman test was used to compare the data of multiple groups. Spearman rank correlation test was used for correlation analysis. P-value of less than 5% was considered to be significant (\**P* < 0.05, \*\**P* < 0.01, \*\*\**P* < 0.001, \*\*\*\**P* < 0.001).

## Limitation of the study

There is one limitation of this study. As we could not obtain enough purified NK cells from peripheral blood of HIV-infected individuals, traditional western blot was not used to detect the protein expression and signaling pathway phosphorylation levels of NK cells. Given that flow cytometry is a well-established and mature technique for analyzing protein and signaling pathway phosphorylation levels, has been widely used, we opted to use flow cytometry to detect relevant indicators.

## Acknowledgments

We thank the volunteers from The First Affiliated Hospital of China Medical University for blood donations. This work was supported by grants from the National Natural Science Foundation of China (82172341), the Mega Projects of National Science Research for the 13th Five-Year Plan (2017ZX10201101), and Scientific research projects of colleges and universities in Liaoning Province (LJKZ0737).

## Author contributions

Y.J. and H.S. conceptualized, supervised the research and revised the manuscript; X.Y. designed research, performed experiments, analyzed data, and prepared the manuscript; J.Z. checked the data; J.L. performed some of the migration experiments; H.G. performed gene expression analyses; Z.Z., Y.F., X.H., Q.H., H.D., and W.G. recruited participants and provided the clinical data.

## Data availability

The published dataset analyzed during the current study was from GEO repository under accession number GSE25669. All relevant data are available from the corresponding author on reasonable request.

## Abbreviations

2-DG: 2-deoxy-D-glucose
2-NBDG: 2-(N-(7-nitrobenzene-2-oxy-1, 3-diazole-4-yl) amino) -2-deoxyglucose)
ADCC: antibody-dependent cell-mediated cytotoxicity
AIDS: Acquired immunodeficiency syndrome
CTL: cytotoxic T lymphocytes
CTLA-4: cytotoxic T-lymphocyte-associated protein 4
CXCL12: C-X-C motif chemokine ligand-12
EAE: experimental autoimmune encephalomyelitis
ECAR: extracellular acidification rate
F-Actin: actin filaments
FBS: fetal bovine serum
GCK: glucokinase
GSEA: Gene set-enrichment analysis
HARRT: highly active antiretroviral therapy
HC: Healthy Control
HIV: human immunodeficiency virus
HIF-1α: Hypoxia-inducible factor-1 alpha
INRs: immune non-responders
IRs: immune responders
KEGG: Kyoto Encyclopedia of Genes and Genomes
LAG-3: lymphocyte-activation gene 3
MFI: mean fluorescence intensity
MCMV: murine cytomegalovirus
NK: Natural killer
SHIP-1: inositol 5-phosphatase-1 containing the SH2 domain
TIGIT: T cell immunoglobulin and ITIM domain

